# Bottlenose whales are more socially connected when in poor body condition

**DOI:** 10.64898/2026.09.25.754507

**Authors:** Sam F. Walmsley, Claire Girard, Katija Newton, David Gaspard, Laura J. Feyrer, Hal Whitehead

## Abstract

Social relationships are strongly linked to health and biological fitness in humans and other animals. For example, social support may buffer individuals against harsh ecological conditions. Reciprocally, better health may allow for greater investment in social relationships. We examined links between health and social behaviour in an endangered population of northern bottlenose whales (*Hyperoodon ampullatus*). Using photo-identifications spanning 37 years, we visually classified whether individuals had poor body condition on each day they were observed (*N =* 3,086 whale-days). We then used generalized linear mixed models to explore how body condition varied with sex, age, and within-individual deviations in group size or other social metrics derived from Bayesian social networks. Approximately 11% of individuals were classified as being in poor condition at any time, though the prevalence of poor condition varied across raters and years. Visually assessed condition was successfully validated with morphometric measures, though there was only weak evidence that poor condition predicted apparent survival. Contrary to expectations, whales in poor condition were found in larger groups and had more social partners than their average across the study period. These findings run counter to the expectation that health and social connectedness are positively correlated in social mammals.

## Introduction

Individual variation in social behaviour has been linked to fitness outcomes across a range of species [1]. Aspects of social integration such as the quality or quantity of one’s relationships can provide direct benefits such as communal defense or information about foraging locations [2], which may ultimately translate into increased rates of reproduction or likelihood of survival. For example, Barbary macaques (*Macaca sylvanus*) with more social partners were more likely to survive the winter, perhaps resulting from increased capacity for social thermoregulation [3]. Of course, being more socially integrated will generally also involve costs such as greater risk of infectious disease or intensified foraging competition [4–6].

Causal links between social behaviour and fitness can be mediated by individual health [1]. Health is a dynamic characteristic of individuals or populations that can be measured across numerous dimensions. Commonly extracted measures of health for animal populations include immunological traits, parasite load, disease status, heart rate variability, and stress hormones [7,8]. There is evidence from a range of non-human species that such measures can be correlated with social connection, though most research thus far has focused on primates. For example, higher-ranking male baboons experience faster rates of wound healing [9], and interaction with preferred partners is associated with reduced urinary stress hormones in chimpanzees (*Pan troglodytes schweinfurthii*; [10]. Despite the general paradigm of health and sociality being positively correlated, the potential for more nuanced (and sometimes negative) associations between specific dimensions of health and sociality may be underestimated. In giraffes (*Giraffa camelopardalis*) for example, infection by gastrointestinal parasites is correlated with interactions between groups [11]. Crucially, such associations will arise from multiple causal pathways – just as social behaviour can determine health outcomes, health status is also known to influence social behaviour. For example, experimental immune challenges lead to reduced social connectedness in vampire bats (*Desmodus rotundus*; [12]) and mice (*Mus musculus domesticus*; [13]).

Body condition reflects an individual’s energy stores and can be one of the most accessible measures of health [14]. While the links between body condition and biological fitness can be complex [15,16], it often correlates with survival or reproductive success in wild populations [17,18]. For example, European badgers (*Meles meles*), with higher body fat reserves have a higher probability of surviving periods of resource scarcity [15]. To the extent that other aspects of poor health can require energetic expenditure (e.g., mounting immune responses, carrying parasites, heightened levels of stress), body condition can be an effective proxy for overall health. In mammals, variation in female body condition can also reflect periods of high maternal investment [19]. Previous research has linked body condition to positions in dominance hierarchies [20,21], or to group size, with larger groups being associated with both higher or lower body condition across studies [22,23]. These associations may also be non-linear: in male prairie voles (*Microtus ochrogaster*), intermediate numbers of social partners tended to have the greatest body mass [24].

Compared to other health metrics (e.g., immunological parameters or heart rate), body condition can be observed from a distance in wild animals, making it an ideal proxy for individual health in inaccessible or at-risk populations where invasive data collection may be impractical or undesirable, such as cetaceans (whales and dolphins). Whales store and metabolize blubber in predictable ways, resulting in distinct visual characteristics of individuals in poor condition such as depressions along the rib cage or behind the melon [25–27]. Poor maternal body condition is associated with reduced offspring growth rates in minke whales (*Balaenoptera acutorostrata*; [28] and humpback whales (*Megaptera novaeangliae*; [29], and has been directly linked to survival in other species – for example, Southern Resident killer whales (*Orcinus orca*) in poor condition have a mortality risk several times higher than other whales in the population [18]. Given the challenge of collecting fine-grain health metrics in wild cetacean populations, body condition provides an important way to measure individual and population health [30].

Using a 37-year individual-based dataset, we tested whether visually assessed body condition correlates with social connectedness in an endangered population of northern bottlenose whales *(Hyperoodon ampullatus).* Northern bottlenose whales are a highly social, deep diving species, forming ephemeral groups of 2-6 individuals [31,32]. Within these fission-fusion societies, individuals maintain multiple long-term preferred social relationships, often with non-kin [32]. Our study aims to address two main overarching questions. First, are there intrinsic or extrinsic factors that predict body condition, such as individual age or sex, or population-level trends over time? Second, is body condition associated with within-individual differences in social behaviour in northern bottlenose whales? In particular, we sought to test the prediction that individuals would be less socially connected when in poor condition, mirroring the general finding that health and fitness are positively correlated with sociality.

## Methods

### Fieldwork and study population

We collected visual and acoustic data of northern bottlenose whales (*Hyperoodon ampullatus*) on the Scotian Shelf during 29 years spanning 1988 to 2025. These whales were primarily found in the Gully submarine canyon (approx. 44° N, 59° W), and occasionally in the adjacent Shortland or Haldimand canyons or other areas of the shelf. We undertook trips to the Scotian Shelf by sailing vessel during the months of June to September, typically for 2-3 weeks at a time. We used a 10m sailing vessel from 1988 to 1992 and a 12m sailing vessel from 1993 to 2025. Data collection methods have varied due to technological advances in photography and other technologies (discussed below), however, the core sampling procedures have remained consistent.

We conducted visual surveys during daylight hours (∼ 0600-2100hrs, depending on the date) throughout the Gully Marine Protected Area (MPA). When we spotted northern bottlenose whales, we recorded date, time, and GPS location. Group size was estimated based on the total number of distinct animals observed within approximately 5 body lengths of one another, using a chain rule, i.e., group members only had to be within the distance threshold to at least one other whale [31]. We photographed the dorsal fin (both left and right) and melon of each individual *ad libitum*. Film cameras were used initially, with digital cameras first being for a small subset of photos in 2002-2003, and exclusively after 2007. For the most recent years of the study (2023–2025), we also collected video footage of the whales from above using an uncrewed aerial vehicle (UAV, or drone; Table S1). See Electronic Supplement for additional detail on the drones used and flight protocols.

### Photographic Analysis

We distinguished and identified individual whales using markings located on the dorsal and dorsal skirt [33]. We used corresponding images of the melon to determine sex based on the relative “roundness”, classifying individuals as female/juvenile or mature male. We determined genetic sex for a small subset of individuals (N=56) using biopsy samples. Calves and juveniles (approximately 1-4 years old) were classified based on visual characteristics including size, colouration, and other characteristics such as fetal folds (See Supplement for additional detail).

### Body Condition Analysis

In cetaceans, body condition can be measured through a range of methods, including anatomical metrics such as blubber mass or lipid content [30]. Morphometric measures of condition from aerial photographs, either from crewed aircraft or (more recently) using drones have become increasingly common (e.g., [34]. Studies have also relied on simple visual classifications by human observers, e.g., for giraffes; [35], as well as cetacean species such as North Atlantic right whales (*Eubalaena glacialis;* [36]), gray whales (*Eschrichtius robustus*; [37], and beluga whales (post-mortem; *Delphinapterus leucas*; [38]. Similarly, a four-level scale ranging from “emaciated” to “robust” was successfully applied to photographs of common dolphins (*Delphinus delphis*) with high agreement across observers [27]. While visual classification lacks the precision of morphometrics, it does not require high-quality aerial video footage, and can potentially be applied to historical photographic datasets to identify long-term trends [37].

We used a historical catalogue of photographs of northern bottlenose dorsal fins and surrounding body parts to assess body condition (Figure 1). Body condition was determined based on four visual indicators: (1) depression around the head/blowhole (similar to the “peanut head” appearance of emaciated killer whales; [25], (2) visible ribs, scapulae, and vertebrae, (3) depression behind the dorsal fin, or (4) depression along the spinal ridge (refer to full protocol in supplement). Such depressions and the visibility of bones generally reflect loss of muscle and/or blubber. Two raters independently scored photographs of whales on each day they were observed as “poor condition” when one or more of the above indicators was identified. We restricted our analysis to left-sided photographs to avoid pseudoreplication as some IDs are known only by a single-side. Results from both analysts were compared and interobserver reliability was quantified with Cohen’s kappa statistic using the *irr* package in R [39,40].

**Figure 1.**
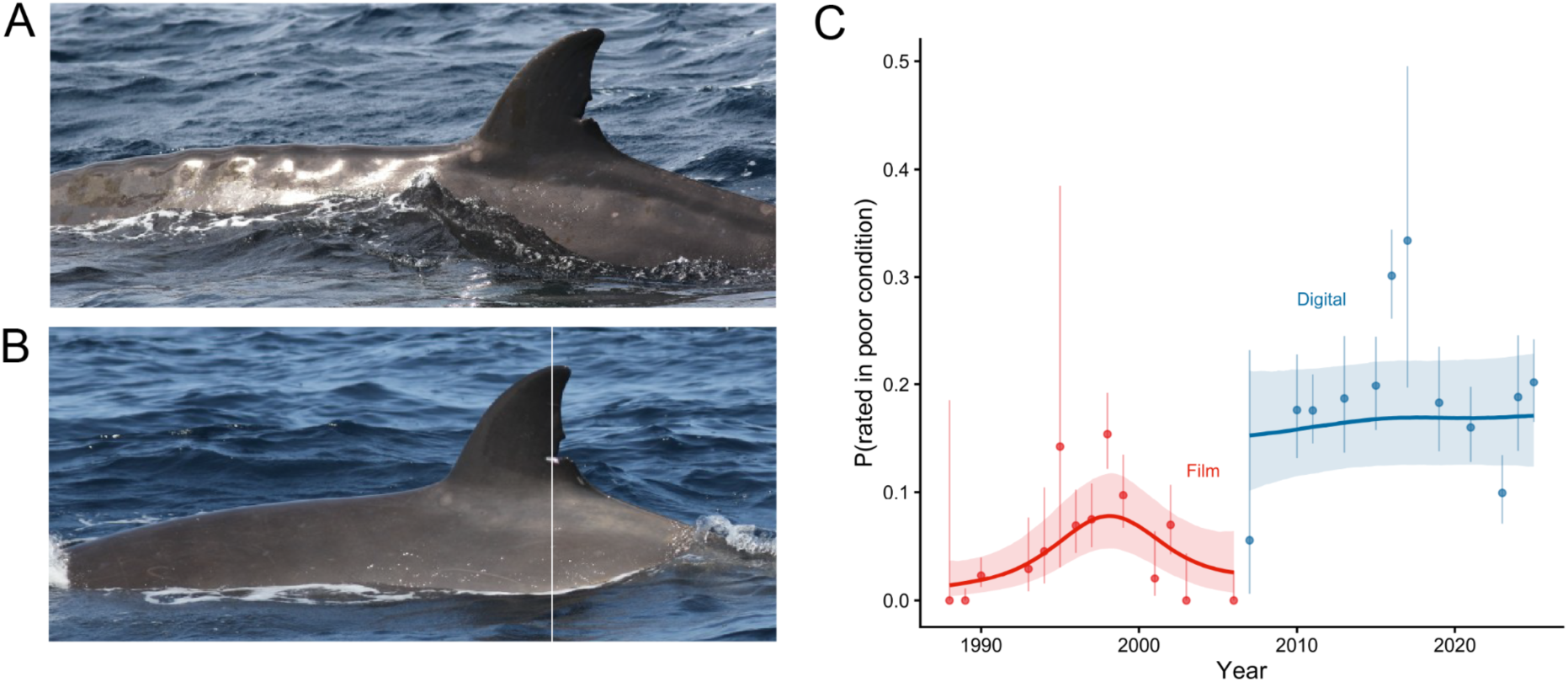
**A)** Example of a northern bottlenose whale observed in poor condition in 2019. **B)** The same whale in improved condition two years later (2021). **C)** Longitudinal trends in poor body condition, estimated using GAMMs while accounting for differences in classification rates for film and digital photography methods. Years with very small amounts of opportunistic data (e.g., < 10 photo-identifications) were excluded for visualization purposes. The curved lines show mean posterior predictions (red: film, blue: digital), with the shaded area showing the associated 90% credible interval. Points represent the annual mean probability of an individual being in poor condition with error bars showing standard errors of the mean.

A subset of these photographic data was used to validate whether visual classifications were correlated with morphometric measures of body condition. Drone footage from 2023-2025 was calibrated and analyzed to estimate a morphometric measure of body condition based on the ratio of body surface area to length (excluding head and tail regions; full details in Supplement). Videos from 2023 were not of adequate resolution to reliably link aerial footage to specific whale IDs. However, we were able to link drone and left-sided photo-ID observations for 45 whales from 2024 and 2025, 7 of which were observed in both years (*N* = 52 whale-years). We used a Bayesian generalized linear mixed effects model (GLMM) with beta-binomial family to test if visual classifications of poor condition were associated with a lower morphometric body condition index (i.e., lower width relative to length). The response variable was the number of days in a given field season (year) when a whale was rated “poor condition”, with the total number of possible ratings for that whale as the number of “trials” (number of days the whale was observed that year). Observations from both raters were included in same model. For predictors we included morphometric body condition, rater ID, and a random effect for each combination of whale ID and year to account for the non-independence of observations from different raters.

### Modelling temporal patterns in body condition

Next, we aimed to characterize long-term trends in body condition over the study period. We used Bayesian generalized additive models (GAMM) with Bernoulli family to relate ratings of poor condition to a potentially non-linear effect of year. Separate smooths were fit for the portion of the study period using film photography and digital photography. We also included fixed effects for photography type, rater ID, and random intercept effects for whale ID and for each combination of whale ID and day. Because this model was based on daily assessments rather than total annual ratings, we also controlled for daily sampling intensity by including the log of the number of photographs per individual (per day) as an additional predictor. We then fit a similar model with a seasonal smooth term (based on Julian day) and random intercepts for years to identify potential seasonal trends in condition. Based on the lack of a clear non-linear effect, we fit an otherwise identical new model with the smooth term replaced by a linear effect of Julian day.

### Modelling links between age, sex, and body condition

Next, we fit a series of GLMMs to explore how body condition varied with 1) sex, 2) age class, and 3) estimated minimum age. Similar to the model linking visual and morphometric condition measures (above), we used the total number of daily “poor” ratings of possible days as a response variable. Each model included a focal predictor of either sex class, age class, or minimum age, as well as controls for photography type, rater ID, and random effects for whale ID, year, and each combination of whale ID and year (to address the non-independence of ratings from multiple observers). With the exception of the model with age class as the main predictor, we excluded observations of individuals classified as young from these models (either calves or juveniles, See supplementary material for additional detail on age classes). We then fit an additional model with only rater ID, photography type, and the same random effects to calculate the proportion of variance explained by individual ID and year, i.e., the “repeatability” of body condition.

### Modelling associations between body condition and probability of survival

If visually-classified body condition in northern bottlenose whales is indicative of a severe detriment to individual health, we would expect it to predict patterns of survival. To test this, we fit Cormack-Jolly-Seber (CJS) mark-recapture models using the *marked* package in R [41]. This approach simultaneously estimates the apparent survival of individuals in subsequent sampling occasions (here, years) alongside the probability of detection or “recapture”. We modelled apparent survival as a function of sex and the proportion of days when an individual was assigned poor condition in a given year, and probability of detection as a function of time (year) and a random effect for individual ID, to account for heterogeneous detection rates across individuals [42]. For years when a whale wasn’t observed, we imputed the most recently observed condition value. We also included an interaction between the type of photography used and the effect of body condition, to test whether ratings made from film vs. digital cameras differed in their usefulness for predicting apparent survival.

### Linking body condition to social connectedness

Similar to [32], we defined social associations between individual whales based on photo-identifications that coincided within 10 minute periods. We then fit a Bayesian social network model to estimate the proportion of time that each pair of whales would be observed together. These measures were calculated for each year. Dyad-specific estimates of the proportion of time spent together (or “edge weights”) within a year were fit using a random effects approach, allowing us to benefit from partial pooling to inform the estimates for pairs of whales with few observations [43]. From these networks we extracted annual measures of degree (number of social partners), eigenvector centrality (a measure of both direct and indirect connections), and mean relationship strength (the mean weight of all edges/connections that an individual formed). Both centrality and mean relationship strength were extracted across 1,000 draws of each annual network, resulting in a full posterior distribution of each value. This allowed us to propagate the uncertainty from the edge weights through to subsequent models.

Hypothetically, body condition could affect sightability or observation rates in a way that could confound certain social measures such as degree and centrality (which would likely be higher for individuals observed more often). For example, individuals in poor condition are likely less buoyant, so raising more of their body out of the water when they breathe and making it easier to take high-quality photographs of their entire dorsal fin. Photographers may also unintentionally take more photographs of animals that are unusually emaciated. We addressed this potential bias using two broad approaches. First, for models based on degree and centrality, we included the number of sampling periods per individual per year as an additional predictor in the model. Second, we also explored the effect of estimated group size on body condition. Group size was estimated during field observations rather than inferred retrospectively from photographic evidence. This provided a measure of social behaviour independent from potential biases of photographic sampling procedures. Field observations of total group size were available for approximately 95% of sightings of northern bottlenose whales. Group size estimates were not available for 2024-2025, so these analyses were based on data from 1988-2023 alone. Photo-identifications of specific individuals were linked to these group size estimates based on time, with a maximum allowable difference of 60 minutes between the recorded time of the photograph and the group sighting (median time difference of 10.8 minutes).

We then fit separate GLMMs to test for associations between body condition and each of these four measures of social connectedness (group size, degree, centrality, and mean relationship strength). We used a within-individual approach whereby both the mean of each individual’s observed social metric across the study period and within-individual deviation in each given year were included as separate predictors. This helped to adjust for potential confounds related to stable differences between individuals [44], and allowed us to ask the more focused question of “*within an individual, are periods of poor condition also periods with more or less social connection?*”.

Again we used a beta-binomial model structure with ratings from both raters included in a single model. We also included a fixed effect for photography type and random intercepts for year, whale ID, and whale-year combination. Here, the main fixed effect of interest was each measure of social connection. We directly incorporated uncertainty in the social measures that were extracted from Bayesian networks (eigenvector centrality and mean relationship strength) using a measurement error approach (see [45]). As described above, controls for the number of sampling periods per year were also included for models with degree and eigenvector centrality. Our aim was to infer overall associations between body condition and sociality, regardless of direction – i.e., our decision to use body condition as the outcome variable does not imply any specific causal interpretation (i.e., that sociality was causing body condition or vice-versa).

Finally, we carried out two robustness checks for these social effects. First fit a series of identical models that included minimum age (both within-and between-individual terms) as a predictor, to assess whether age could be driving within-individual correlations between sociality and condition. Second, we fit a similar model with a quadratic effect of each social measure, to test for non-linear trends such as enhanced body condition at intermediate group sizes. For these quadratic models we did not decompose within-individual from between-individual patterns, and were unable to incorporate measurement error in eigenvector centrality and mean relationship strength.

All GAMMs and GLMMs, including the social network model, were fit using the *brms* package in R with weakly informative priors [46]. Effects from frequentist mark-recapture models are reported as CIs and 95% confidence intervals. Unless otherwise stated, all other effects are reported as posterior means, 90% credible intervals, and probabilities of directional effects (*p*_dir_).

## Results

We examined 23,529 left-sided photographs to assess body condition across the long-term dataset northern bottlenose whale photo-identifications. This included observations from 743 individual whales from 29 years of field observations, of which 112 individuals were classified as male, 262 as female-juvenile, and 369 with unclear sex. The frequency with which individuals were classified as having poor body condition varied between observers. Of 3,086 whale-day combinations, 5.8% and 21.3% were classified as being in poor condition by observers 1 and 2, respectively (overall mean 13.6%). Of individuals seen on a given day, 11% were assigned poor condition on average. Classifications from different raters showed “fair” but statistically significant agreement (Kappa = 0.255, p < 0.001). There was slightly higher agreement during years when exclusively film cameras were used (Kappa = 0.339, p < 0.001) compared to more recent years (Kappa = 0.214, p < 0.001), though there was significant reliability within each period.

Whale ID explained most of the variation in condition after accounting for rater ID and photography type (repeatability on logit scale R = 0.71, CI [0.58, 0.83]), with year playing a smaller but still detectable role (R = 0.12, CI [0.05, 0.23]). We found strong evidence for a non-linear trend in body condition during the early portion of the study period, with worsening condition in the late 1990s (Figure 1C). We did not detect any obvious inter-year trends in the more recent period when digital cameras were used, but there was still substantial variation across years – for example, there was a particularly high prevalence of poor condition in 2016 (Figure 1). We found no strong evidence for a non-linear effect of season (Figure S2), but detected a slight linear effect of Julian day on the probability of whales being assigned poor condition (*β* = 0.01, 90% CI [3.7 × 10^-3^, 0.02], *p*_dir_ = 99.4%).

As predicted, individuals in better morphometric condition were less likely to be assigned “poor condition” by visual observers (*β* = -0.6, 90% CI [-1.2, -0.07], *p*_dir_ = 96.7%). Still, there was non-trivial variability in morphometric measures among individuals assigned “poor” (Figure 2A), suggesting that visual classifications were a valid, but coarse, proxy of morphometric condition.

**Figure 2.**
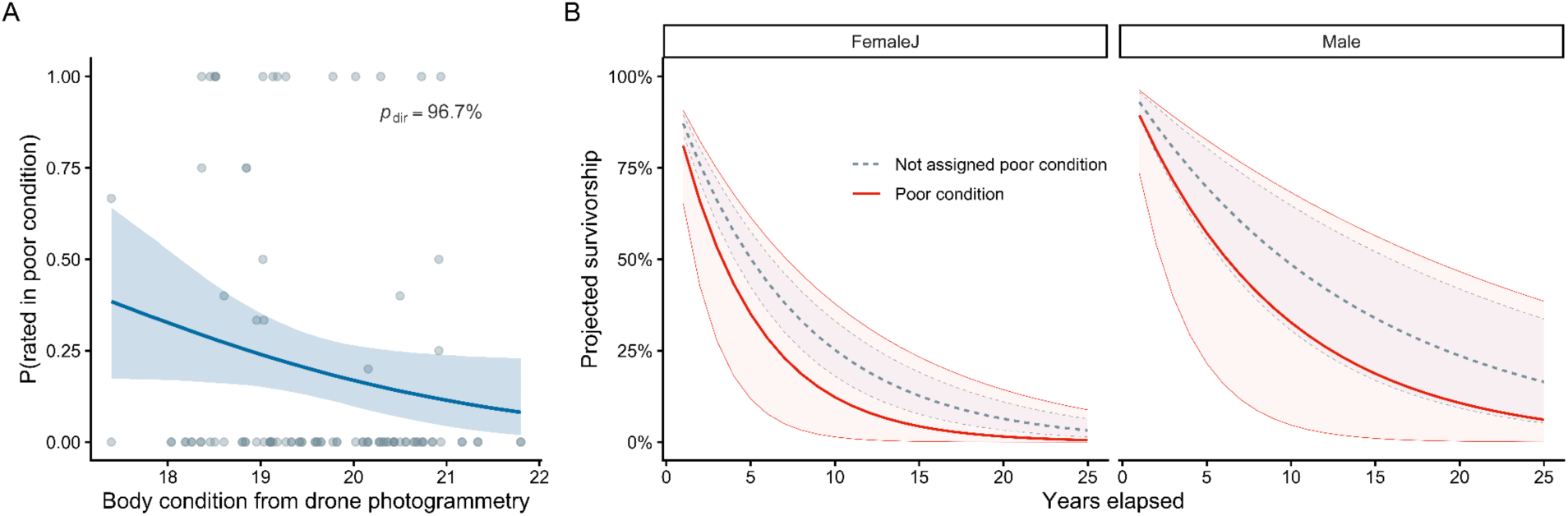
A) Association between morphometric estimates of body condition and visual classifications of poor condition from data collected between 2024-2025. Higher values of body condition indicate greater width relative to length, assumed to reflect enhanced energy reserves. B) Apparent survival of northern bottlenose whales as a function of sex classification (left: female-juvenile, right: male) and visually classified body condition (grey = 0% poor condition ratings, red = 100% poor condition ratings), showing effects calculated for the “digital” era of the long-term study.

### Associations between body condition and sex or minimum age

Sex and age were also associated with body condition. Male northern bottlenose whales were approximately twice as likely to be assigned poor body condition than female-juveniles (*β* = 0.85, 90% CI [0.41, 1.31], *p*_dir_ = 100%; Figure S1). We also found that estimated (minimum) age was associated with poor body condition (*β* = 0.09, 90% CI [0.06, 0.11], *p*_dir_ = 100%), consistent with declining physical health in ageing whales. However, we detected no clear difference in the probability of calves or juveniles being assigned poor condition, relative to adults (calves: *β* = -0.79, 90% CI [-2.32, 0.67], *p*_dir_ = 82%; juveniles: *β* = -0.17, 90% CI [-0.80, 0.43], *p*_dir_ = 67.2%; Figure S1).

### Associations between body condition and survival

We used sighting histories to estimate the effect of poor body condition on apparent survival. Poor condition was negatively associated with apparent survival when digital photography was used, though the effect was quite uncertain (*β* = -0.46, 95% CI [-1.05, 0.13], Figure 2B). The association between apparent survival and condition appeared to be stronger in digital years, though we found no strong evidence for a statistical difference between effects (interaction term between condition and photography type: *β* = 1.04, 95% CI [-0.19, 2.28]). Sex played a stronger role in apparent survival, with males having higher apparent survival compared to female-juveniles (*β* = 0.68, 95% CI [0.39, 0.96]). Individuals of unknown sex had lower apparent survival (*β* = -1.28, 95% CI [-1.53, -1.04]), which may reflect differences in residency patterns (e.g., individuals that use the Gully less would be more likely to “disappear” and would also lack sex data, by virtue of being observed fewer times). Apparent survival was low overall in this model, resulting in average year-to-year apparent mortality estimates of 13.5% for female-juveniles and 7.5% for males. Apparent survival was much lower for individuals of unknown sex (mortality of 35.6%), suggesting that movement or residency patterns are likely important.

### Associations between body condition and social connectedness

Contrary to expectations, whales in poor condition had similar and sometimes stronger measures of social connection than they did at other times in the study period (Figure 3). More specifically, poor condition was associated with within-individual increases in both group size (*β* = 0.13, 90% CI [0.05, 0.20], *p*_dir_ = 100%), and number of social partners (adjusted for sampling intensity; (*β* = 0.04, 90% CI [9.10 × 10^-3^, 0.06], *p*_dir_ = 99%). In other words, whales in poor condition had more social associations compared to periods of their lives when they were in better condition. There was no clear evidence of an increase in eigenvector centrality (*β* = 0.55, 90% CI [-1.10, 2.20], *p*_dir_ = 71%), and no evidence of a difference in mean relationship strength (*β* = -2.50 × 10^-3^, 90% CI [-2.46, 2.47], *p*_dir_ = 50%). Over and above within-individual changes, we found no indication that *between*-individual differences in sociality were associated with the prevalence of poor condition (lower panels in Figure 3). Similar within-individual associations between social traits and body condition were found when conditioning on minimum age. Consistent with the overall trend between condition and age, these models also pointed to within-individual ageing as an independent predictor of poor condition (See Figure S3). We also found little evidence for non-linear effects – while models with quadratic effects suggested declines in body condition at extremely high values of group size and number of partners, peak body condition for each occurred above the 95th percentile of the social trait (Figure S4).

**Figure 3.**
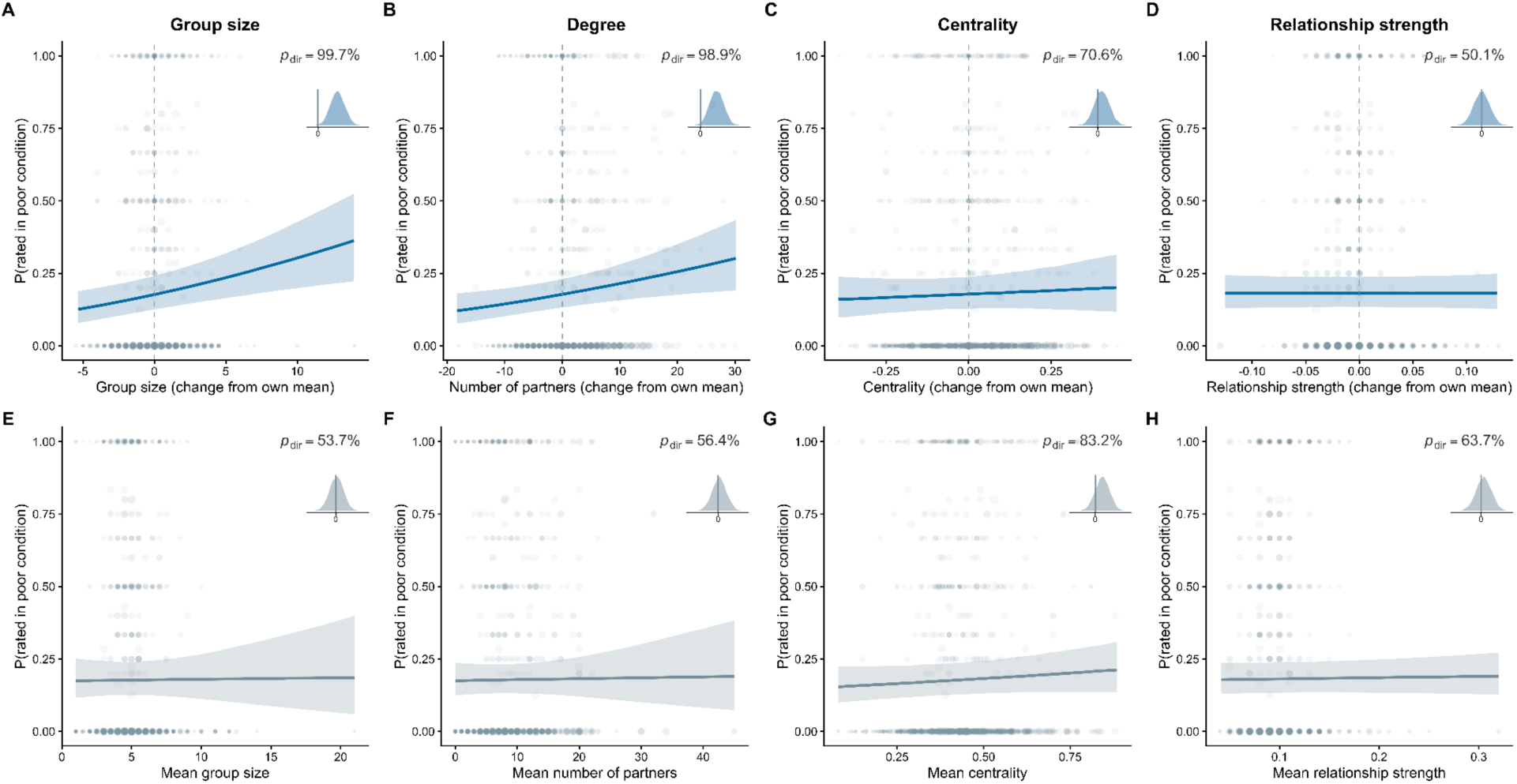
Associations between visually-assessed poor body condition and social connection in northern bottlenose whales. Grey points represent raw data (binned). Blue lines represent the mean posterior prediction of within-individual effects (A-D), with the blue ribbon showing the corresponding 90% credible interval, while grey lines and ribbons (E-H) represent between-individual effects. Smaller inset plots show the full posterior of the association of body condition and each measure of social connection, alongside the posterior probability of a directional effect.

For each of these social models, photography type and rater ID had very similar effects on the prevalence of poor condition: there was a lower probability of poor condition with film photography, and an increased probability of poor condition with the second rater. For the models that conditioned on sampling intensity, the number of sampling periods by year had no clear effect on condition ratings (degree model: *β* = 4.70 × 10^-3^, 90% CI [-0.05, 0.06], *p*_dir_ = 55%; eigenvector centrality model: *β* = 0.03, 90% CI [-0.01, 0.08], *p*_dir_ = 88%).

## Discussion

Evidence from many species points to positive correlations between the health and social connectedness of individuals [1]. Focusing on a small population of northern bottlenose whales, we found the opposite: whales in poor body condition were found in larger groups and had more social partners relative to their own average across the study period. This finding emphasizes that while links between social behaviour and health are likely widespread, they may be highly variable across populations or dimensions of health or social integration.

There are several causal pathways that could contribute to this unexpected association between health and sociality in northern bottlenose whales. First, changes in body condition could cause differences in social behaviour. This could involve greater attraction to conspecifics when in poor health – for example, prairie voles (*Microtus ochrogaster*) showed greater motivation to affiliate with their preferred partners when subjected to an immune challenge [47]. This relates to theory on “sickness behaviour”, i.e. changes in behaviour associated with illness [48]. While behavioural changes can be by-products of health status (e.g., reduced energy resulting in less movement), they may also confer adaptive benefits to the host or the host’s social partners [49]. For example, grey wolves (*Canis lupus*) infected by sarcoptic mange have better chances of survival if they live in larger packs, particularly when packmates were not infected themselves [50]. Northern bottlenose whales in poor condition might be drawn to larger groups if they are in greater need of potential benefits like social foraging (see [51]), or protection from predators. Interestingly, we observed no change in mean relationship strength, implying that the increase in group size and number of partners may have resulted from an increase in total social effort, rather than a prioritization of more (but weaker) dyadic associations. Health-induced changes in association could also be driven by social partners, e.g., via caregiving, which has been documented in several species (often eusocial insects or mammals; [52]). A caregiving explanation would be consistent with early reports that northern bottlenose whales would gather and remain with individuals injured by whalers until they died [53]. More broadly, epimeletic behaviour (tending to sick, injured, or deceased conspecifics) is prevalent in cetaceans [54]. Some form of care-giving would also be consistent with evidence of long-term social preferences in the population, particularly within the sexes [32]. These stable affiliations may simply be resilient to fluctuations in condition, especially given that poor condition (as defined in this study) can be transient (Figure 1). Attraction to females with young offspring could also contribute to this pattern [55], if nursing reduces body condition, though this is unlikely to be the dominant mechanism given that females were less likely to be in poor condition than males.

The reverse causal pathway may also contribute to these results, i.e., variation in social behaviour could be causing changes in body condition. For example, whales spending time in larger groups during a given year may experience greater foraging competition, or a higher burden of parasites or disease – group size and eigenvector centrality are commonly used proxies for the risk of infectious disease [56,57]. Classically, such effects are expected to contribute to a pattern of optimal group size where downstream consequences for physiology and survival are maximized, with smaller and larger groups being more costly [58]. However, links between body condition and social connectedness appeared to be mostly linear in our study, with evidence of potential declines in condition only at unusually high levels of sociality. More fine-grained longitudinal data on changes in both social behaviour and body condition (i.e., throughout the year) could help to resolve which of these causal pathways is most important [59]. Our within-individual approach controlled for any stable unobserved confounds at the level of individual whales, and our robustness checks based on estimated (minimum) age suggested that age was not confounding the observed health-sociality correlations. Still, it is possible that other time-varying confounds such as reproductive state are causing coincident changes in sociality and health within individuals over time.

Beyond these links to social connectedness, body condition in northern bottlenose whales was also directly associated with sex and minimum age. If variation in condition were mainly driven by female reproductive investment, we would expect females to have a higher prevalence of poor condition overall. Instead, the finding that males were worse off may point to other factors, such as sex-specific resource availability, foraging dynamics, or movement patterns being responsible. Diet differences between the sexes [60], for instance, could result in differential vulnerability to variation in prey availability and quality. One limitation of our study is our inability to track the whales throughout the year, raising the possibility that sex differences in body condition could reflect the higher energy investment of males during the mating season, which runs from late July to August and overlaps with data collection [61]. Male headbutting, observed throughout the study, is likely a form of male-male competition [62] and parallels head-to-head contests in terrestrial ungulates [63]. Males also typically exhibit weaker immune systems than females [64], raising the (speculative) possibility that infectious disease or parasite loads could be shaping sex differences in condition. Interestingly however, males in our study were also found to have higher rates of apparent survival overall, confirming a prediction based on the age structure of whaled specimens that male northern bottlenose whales live longer than females [65]. While we had a limited capacity to distinguish individual age from temporal trends (e.g., Figure 1), the finding that whales known to be older were more likely to be in poor condition is consistent with senescence in wild populations [66].

We found weak evidence that a simple measure of body condition extracted from photographic data was associated with apparent survival. Heightened mortality risk among individuals in poor condition has been observed in other cetaceans [18,36,67], suggesting that body condition may be useful as a rapid assessment tool for population health and viability. In this case, we found substantial variation in the prevalence of poor body condition across the 37-year study period, though there has been relative stability over the past decade. Like other beaked whales, this population faces a range of ongoing threats including injury or entanglement by fisheries gear as well as the accumulation of ocean plastics, which has been linked to starvation [68,69]. However, this particular population has been undergoing a rebound in abundance since the implementation of a Marine Protected Area in 2004 [70]. Beyond the direct effects of anthropogenic activities, factors such as prey abundance or climatic variation are also expected to influence cetacean body condition over time [71,72], and may play a role in the trends we observed (Figure 1).

Our initial expectation was that the widespread health benefits of social integration and the greater capacity for healthier individuals to invest in social relationships should lead to a positive correlation between body condition and social metrics in northern bottlenose whales. Instead, we found that within individuals, periods of poor condition coincided with increases in group size and number of social partners. These findings suggest that links between health and sociality are complex and unlikely to be homogenous across ecological contexts.

## Supplemental Material

**Figure S1.**
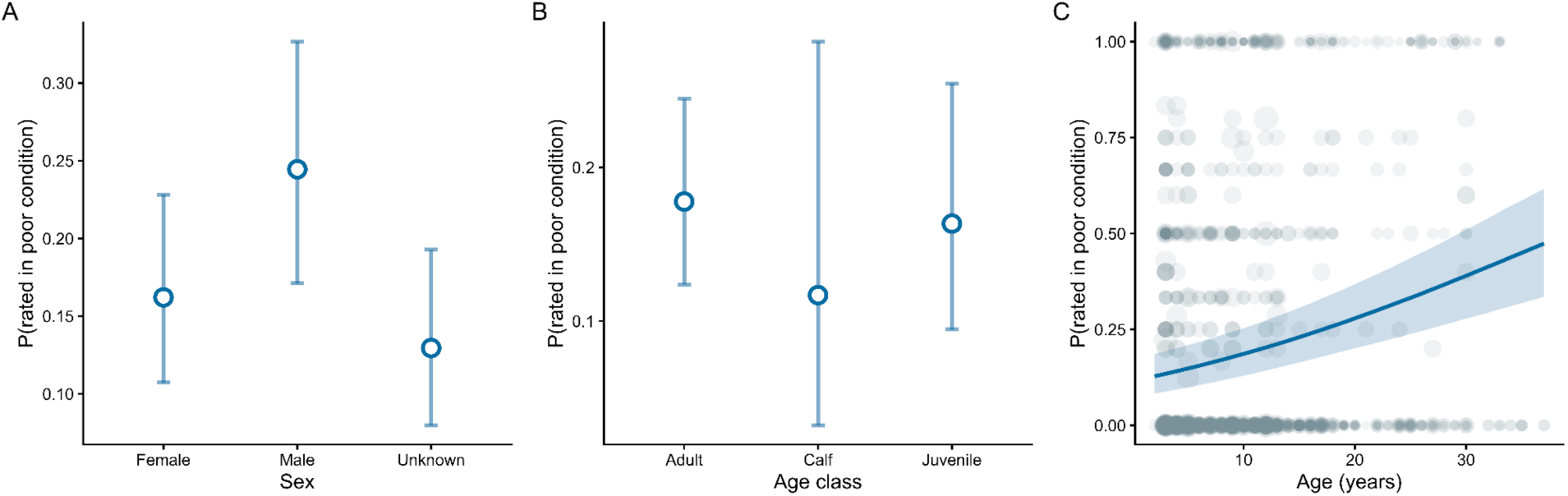
**A)** Predicted association between sex and poor condition. **B)** Predicted association between age class and poor condition. **C)** Predicted association between estimated (minimum) age and poor condition. Grey points represent raw data (binned), with the size of the point representing the number of observations per bin.

**Figure S2.**
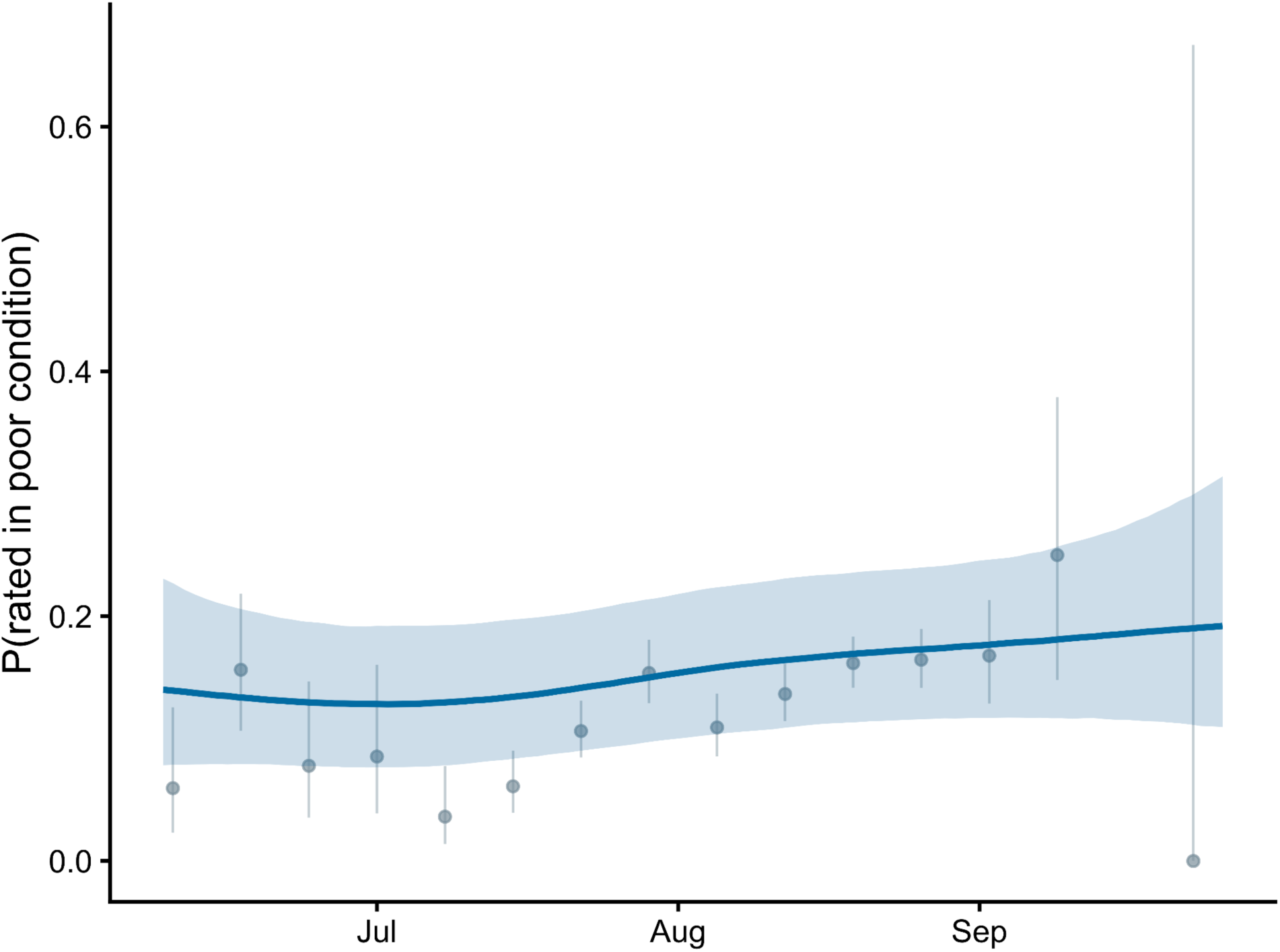
Estimated seasonal trend in body condition. The blue line shows the predicted mean probability of a “poor condition” rating as a function of Julian day while the shaded portion shows the associated 90% CI. Points represent raw weekly means pooled across raters, with associated 95% intervals.

**Figure S3.**
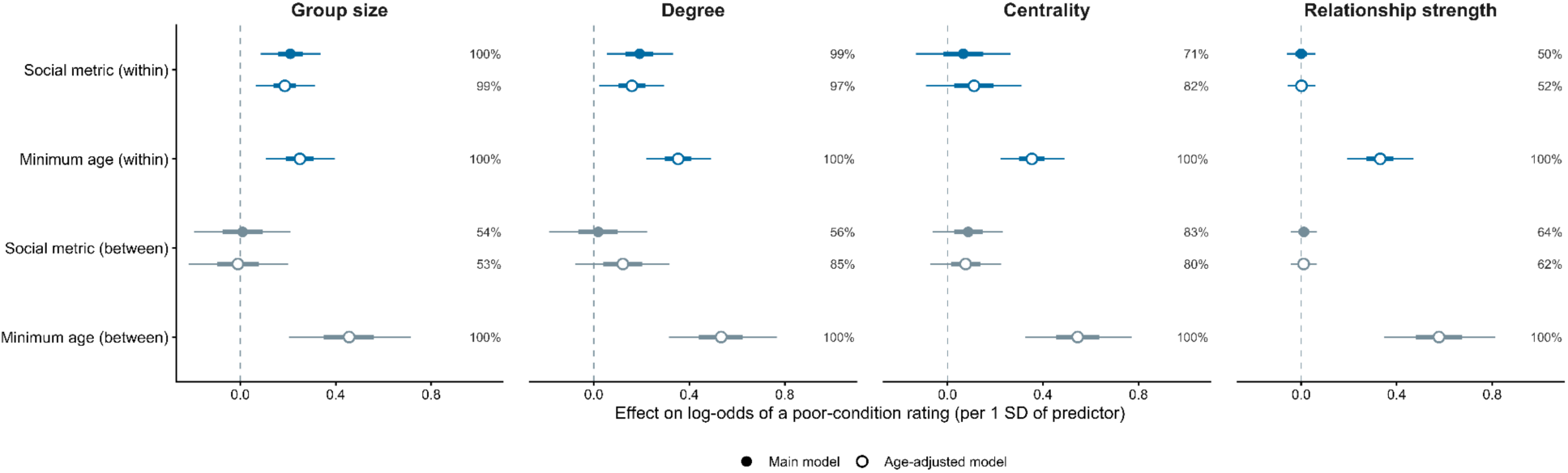
Effects of within-individual deviations in social behaviour on body condition are similar when accounting for estimated (minimum) age. Points show posterior means (filled for model excluding minimum age), while thick and thin portions of the horizontal bars represent 50% and 90% CIs, respectively.

**Figure S4.**
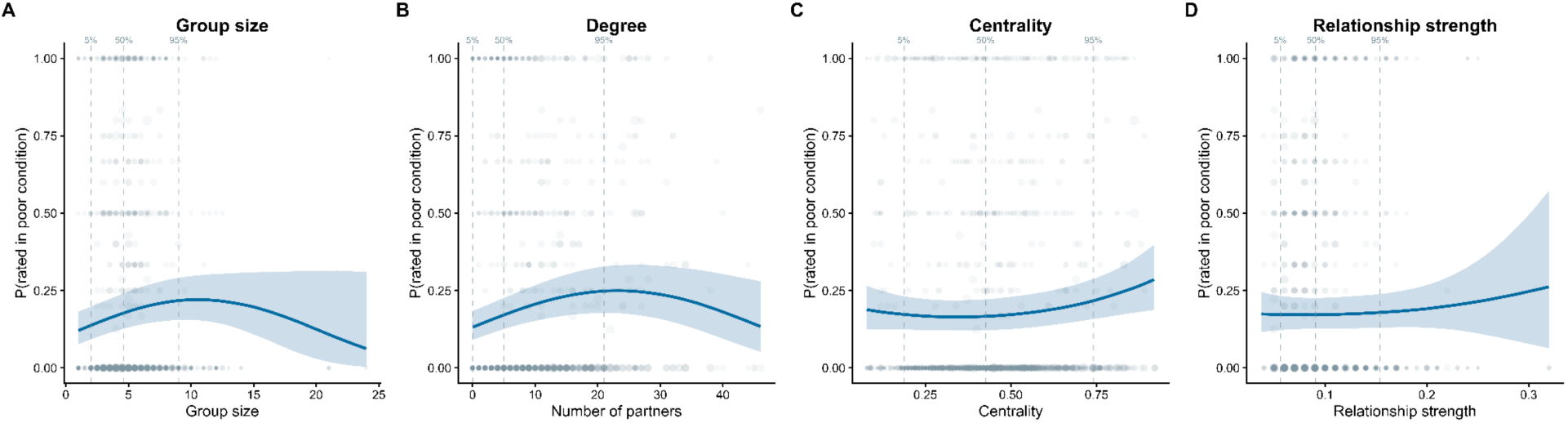
Associations between social behaviour and body condition using non-linear (quadratic) effects. Vertical lines show the 5th, 50th, and 95th percentiles of each social trait to help visualize the distribution of social trait observations. While we find evidence of declining body condition at very high values of group size or number of social partners, these relationships were effectively linear across the large majority of variation in social traits.

### Age classifications

Calves were identified through size, presence of an angled dorsal fin, and fetal folds along the body. Calves have grey bodies with a distinct black stripe along their spine beginning just behind their dorsal fin. Their melons are generally lighter in colour and sometimes have a “wrinkled” appearance. Their beaks were also distinctly smaller in proportion to their melon in comparison with adults. The identification of juveniles can be more ambiguous as they lack distinctive characteristics such as fetal folds. Juveniles were identified as being larger than calves and smaller than adults (up to ⅔ the size of an adult whale). Juveniles were also visually identified by a darker band along the spinal ridge following the dorsal and an absence of scarring compared to adults. The main characteristics used to consistently identify juveniles were a comparatively small dorsal fin and brown body colour.

### Minimum age

Following [1,2], we calculated minimum age based on the number of years since an individual was first observed in the long-term photo-identification catalogue. We added 1 year for animals first seen as juveniles (who would be at least 1 year old), and added 3 years for animals that did not appear to be calves or juveniles.

### Protocol for classifying body condition from photo-identification data

Images of identified individuals from the finalized dorsal and melon photo ID catalogue were evaluated to determine body condition. Individual whales were identified through reliable markings located on the dorsal and dorsal skirt. Body condition was determined based on four visual indicators: depression around the head/blowhole (similar to the “peanut head” appearance of emaciated killer whales; [3], visible ribs, scapulae, and vertebrae, depression behind the dorsal fin, and depression along the spinal ridge. Generally, these depressions and the visibility of bones are expected to reflect the loss of muscle and/or blubber. Two analysts independently scored the images from the full catalogue of dorsal photographs as “poor condition” when one or more of the above indicators was identified.

### Aerial video collection

Drones were hand-launched from the sailing vessel during whale encounters, when weather was at or below Beaufort Wind Force 3. The drones were equipped with a LiDAR system sampling at 1 Hz. The LiDAR measured the vertical distance between the laser and the sea surface, which was subsequently corrected for drone pitch and roll [4] and LiDAR position relative to the camera. The corrected LiDAR altitude was used for photogrammetric scaling of morphometric measurements (i.e., to convert pixel-based measurements from aerial images into absolute body dimensions) [5].

Prior to take off, video recording started and a device displaying GPS time was placed in front of the camera. This was to facilitate the alignment of the time stamp between the video time and LiDAR altitude recordings [6]. Video recording was stopped after landing and before powering down the drone. Videos of whales were recorded instead of stills to increase likelihood of imaging whales in optimal position.

All videos were recorded from altitudes > 15 m (mean = 18.5 m) with the camera at nadir (pointing 90° down), aiming to position whales lengthwise and centred in the frame to maximise resolution. Each whale within a flight or group was systematically centered in the frame for several surfacings to ensure sufficient images to extract morphometric measurements would be collected for each individual. Flight duration averaged 12 minutes. Up to three flights per group were conducted if whale behaviour did not allow all individuals to be properly imaged in a single flight.

**Table S1.**
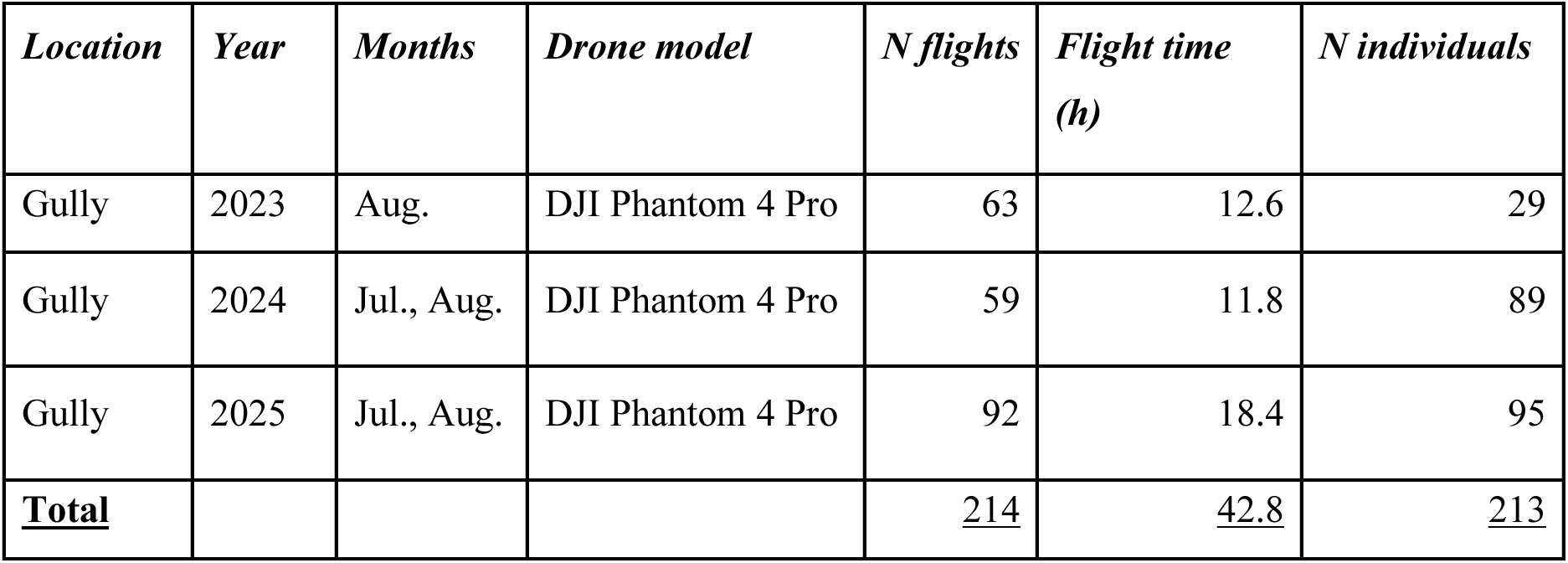
Summary of drone survey effort for northern bottlenose whales in the Gully by year, including number of flights, total flight time, and number of individuals measured from aerial images. Note that not all individuals in aerial videos were able to be confidently linked to a unique ID from the long-term study.

| *Location* | *Year* | *Months* | *Drone model* | *N flights* | *Flight time*<br>*(h)* | *N individuals* |
| --- | --- | --- | --- | --- | --- | --- |
| Gully | 2023 | Aug. | DJI Phantom 4 Pro | 63 | 12.6 | 29 |
| Gully | 2024 | Jul., Aug. | DJI Phantom 4 Pro | 59 | 11.8 | 89 |
| Gully | 2025 | Jul., Aug. | DJI Phantom 4 Pro | 92 | 18.4 | 95 |
| **<u>Total</u>** |  |  |  | <u>214</u> | <u>42.8</u> | <u>213</u> |

### Protocol for estimating morphometric body condition from aerial video

Still images were then systematically extracted for all whales visible in the videos, using VLC software. Still images timestamps were recorded during extraction from videos to align each image with the corresponding altitude. Images were selected when individuals were positioned as close to horizontal as possible at or near the surface, with body length and width clearly visible. Up to five suitable still images per individual were extracted to account for variation due to changes in body position and measurement error [4,7]. Each image was quality rated to assess its suitability for morphometric measurements. Images rated as poor quality were discarded. Retained still images were corrected for lens distortion prior morphometric measurements using the Computer Vision Toolbox in MATLAB [7].

Morphometric measurements were manually extracted from the still images using the Morphometrix V2 software [8]. Measurements included total body length and width measured at every 5% increments of body length, for a total of 20 width measurements. The body surface area was calculated as a series of trapezoids, with the outer edge of each segment approximated as a parabola, to estimate the flat surface area of the body for each width segment [7]. Then, body condition was calculated as the ratio of body surface area between the 15% and 80% width measurements to body length. Restricting our analysis to the 15% - 80% portion of the body allowed us to focus the analysis on body areas that are the most likely to fluctuate in size and shape according to nutritional status, as the head and tail regions are unlikely to be areas of significant lipid stores (see [9]).

A total of 72 unique individuals imaged by drone were linked to the dorsal-fin catalogue and assigned a dorsal ID, representing 81 individual-year records (28 in 2024 and 53 in 2025), with nine individuals resighted in both years. Following image quality control, 210 still frames were retained for morphometric measurements and drone-derived body condition estimates, with a median of two frames per individual-year (range: 1–8). Repeated measurements within each individual-year were collapsed to a single value by taking the median for each morphometric measurement, including total body length and body-width measurements at standardized positions along the body, before calculating body area index. Multiple retained frames could originate from the same sampling date or from repeated encounters across the field season. Of the 81 individual-year records, 66 (81.5%) were sampled on a single date, 14 (17.3%) on two dates, and one (1.2%) on three dates. Among the 15 individual-year records sampled on multiple dates, the interval between the first and last measurement ranged from 1 to 25 days (median = 24 days; mean = 15.7 days).

